# Robust single-cell DNA methylome profiling with snmC-seq2

**DOI:** 10.1101/294355

**Authors:** Chongyuan Luo, Angeline Rivkin, Jingtian Zhou, Justin P. Sandoval, Laurie Kurihara, Jacinta Lucero, Rosa Castanon, Joseph R. Nery, António Pinto-Duarte, Brian Bui, Conor Fitzpatrick, Carolyn O’Connor, Seth Ruga, Marc E. van Eden, David A. Davis, Deborah C. Mash, M.Margarita Behrens, Joseph R. Ecker

## Abstract

Single-cell DNA methylome profiling has enabled the study of epigenomic heterogeneity in complex tissues and during cellular reprogramming. However, broader applications of the method have been impeded by the modest quality of sequencing libraries. Here we report snmC-seq2, which provides improved read mapping, reduced artifactual reads, enhanced throughput, as well as increased library complexity and coverage uniformity compared to snmC-seq. snmC-seq2 is an efficient strategy suited for large scale single-cell epigenomic studies.

---

Several library preparation strategies for single-cell DNA methylome profiling have been developed based upon the Post-Bisulfite adapter Tagging (PBAT) approach and have been applied to study epigenomic heterogeneity in embryonic stem cells, mouse and human brains, and preimplantation embryos ^1–4^. Most library preparation methods for single-cell methylomes use random-primed DNA synthesis to incorporate the upstream sequencing adapter but vary in the strategies that incorporate the downstream adapter sequence. A second round of random-primed synthesis is used by scBS-seq whereas sc-WGBS and snmC-seq use proprietary 3’-adaptor tagging methods ^2,3,5^. A combinatorial indexing strategy has recently been applied to generate single-cell methylome libraries without physically compartmentalizing individual cells ^6^. Sequencing libraries generated by existing single-cell methylome methods suffer from one or more of the following problems: high levels of artifactual sequences such as adapter dimers, low mapping rate, or small insert size. We previously developed snmC-seq, which provides improved reads mapping ^5^. However, compared to traditional MethylC-seq ^7^, snmC-seq libraries still contain substantially higher levels of adapter dimer sequences and have lower read mapping rates. We systematically examined experimental factors that can be modified to improve the quality of snmC-seq libraries and resulted in the development of snmC-seq2 (Fig.1a). A detailed step-by-step bench protocol is provided in Supplementary File 1.

**Figure 1.**
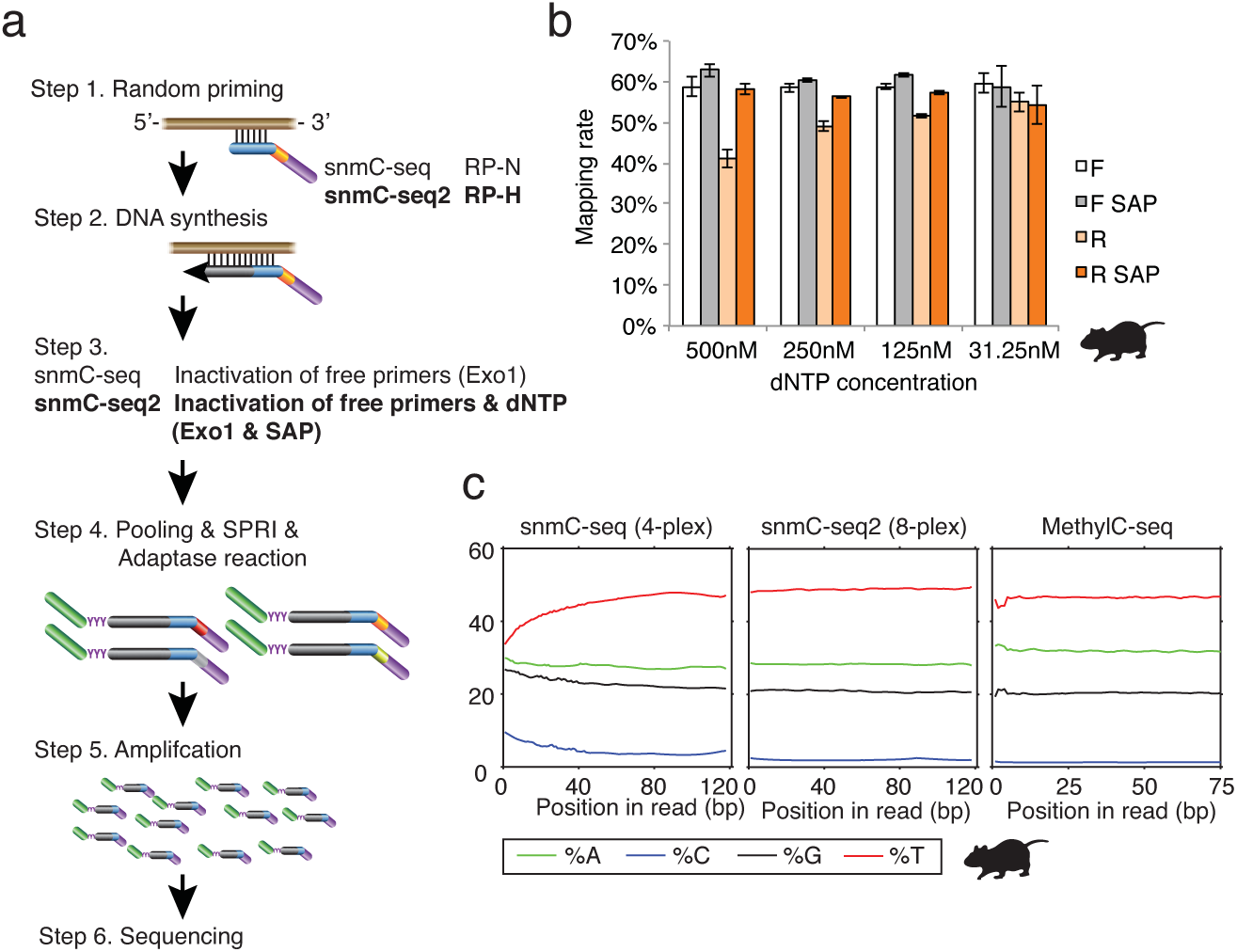
Improvement of single-cell methylome reads mapping. (**a**) Schematics of snmC-seq workflow. Steps modified in snmC-seq2 are highlighted. RP-N and RP-H indicate random primers used in snmC-seq and snmC-seq2, respectively. Exo1 - Exonuclease 1. SAP - Shrimp Alkaline Phosphatase. SPRI - Solid Phase Reversible Immobilization. (**b**) Reverse reads (R) mapping is improved by reducing nucleotide concentration in random-primed DNA synthesis reaction or treatment with shrimp alkaline phosphatase, whereas Forward reads (F) mapping is not affected. (**c**) snmC-seq2 produces reverse reads with comparable base composition as MethylC-seq reads.

We first set out to develop strategies to improve the mapping rate of snmC-seq reads by examining the snmC-seq dataset generated from mouse frontal cortical neurons ^5^. We found that the mapping rate of reverse reads (R) generated by paired-end sequencing is significantly lower in multiplexed (4-plex) than non-multiplexed (1-plex) snmC-seq (Supplementary Fig.1a, p=1.6×10^−32^, t-test)^5^, whereas the mapping rate of forward reads (F) was comparable between multiplexed and non-multiplexed libraries. We identified an aberrant base composition in the reverse reads of multiplexed snmC-seq data (Supplementary Fig.1b). After the conversion of >99% unmethylated cytosines (C) to uracils (U) with bisulfite treatment, only 1.2% nucleotides of mouse frontal cortical neuron genomes are expected to be read as C during sequencing ^8^. However, multiplexed snmC-seq data showed an elevated frequency of C at the beginning of reverse reads and reached a maximum of 9.6% (Supplementary Fig.1b). We hypothesized that the aberrant elevation of C frequency was caused by carryover nucleotides triphosphate (dNTP) that were incompletely removed by DNA purification using Solid Phase Reversible Immobilization (SPRI) beads after the random-primed DNA synthesis (Fig. 1a). The subsequent 3’-tagging by Adaptase involves simultaneous tailing and adaptor ligation (Step 4 in Fig. 1a). While random-primed DNA synthesis requires all four nucleotides (A, T, C and G), the tailing reaction mediated by Adaptase uses only two nucleotides and generates a synthetic low-complexity, short tail sequence with an average length of 8 bp that is present at the beginning of reverse reads. Contamination of the Adaptase reaction by a dNTP mixture leads to the formation of synthetic tail sequences of high-complexity base composition to the 3’-end of the random primed DNA synthesis products. Since this tailing reaction can extend to up to 50 bp (Supplementary Fig.1b), the long synthetic tail sequences were not sufficiently trimmed by snmC-seq trimming condition (16 bp from both 5’ and 3’) and caused mapping failure for reverse reads that retained synthetic tail sequences. We tested this nucleotide carryover hypothesis by supplying different amounts of dNTP (31.25 to 500 nM) to the random-primed DNA synthesis reaction (Fig.1b). Reducing dNTP concentration effectively alleviated the aberrant elevation of C frequency at the beginning of reverse reads (Supplementary Fig.1c), and increased the mapping rate of reverse reads (Fig.1b). To develop a procedure that robustly prevents dNTP contamination, we incorporated into snmC-seq2 a dephosphorylation step before the Adaptase reaction using a temperature sensitive Shrimp Alkaline Phosphatase (SAP) to inactivate nucleotide triphosphates (Fig.1a). The SAP treatment completely suppressed the aberrant base composition observed in the reverse reads generated by snmC-seq (Fig.1c). We incorporated the SAP treatment to snmC-seq library preparation (snmC-seq + SAP) from human frontal cortex (Brodmann area 10, BA10) and validated that the dephosphorylation of dNTP significantly improved the mapping of reverse reads (Supplementary Fig.1d and Supplementary Table 1, p=2.4×10^−58^, t-test). We found that the addition of SAP treatment had no impact on the sequencing library complexity (Supplementary Fig.1e).

**Table 1.**
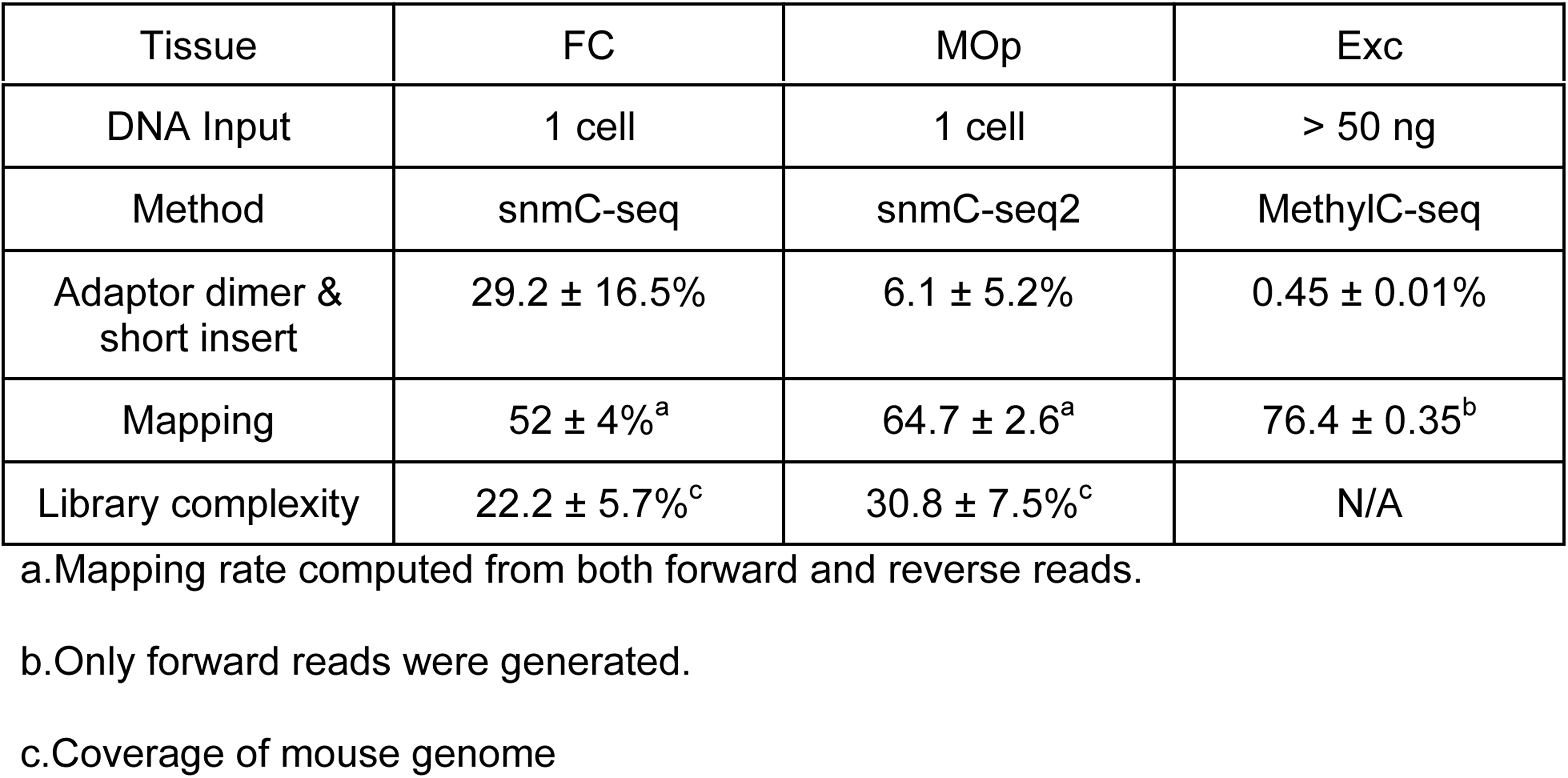
Comparison of the quality of methylome libraries generated by snmC-seq, snmC-seq2 and MethylC-seq. FC - mouse frontal cortex. MOp - mouse primary motor cortex. Exc - Camk2a+ expressing mouse cortical excitatory neurons.

Sequencing reads containing adapter dimer sequences are a major type of artifact in snmC-seq libraries ^5^. Indeed, libraries generated from human medial frontal gyrus using snmC-seq contained an average of 22.6±9.5% adapter dimer sequence or short insert (Fig.2b). The main source of such artifacts is carryover of unused random primer into the Adaptase reaction. We hypothesized that some unused random primers are present in partially double-stranded form caused by primer-to-primer hybridization and are resistant to Exonuclease 1 digestion (Step 3 in Fig.1a). We speculated that adapter dimers could be reduced by removing the guanine (G) base in the degenerated 3’ arm of snmC-seq random primers (Fig.2a). Random primers used in snmC-seq and snmC-seq2 are hereafter referred to as RP-N (N=A,T,C,G) and RP-H (H=A,T,C), respectively. The use of RP-H was expected to destabilize primer-to-primer hybridization by preventing the formation of more stable G:C pairs. We also expected that the hybridization between RP-H and bisulfite converted genomic DNA to be minimally affected, since greater than 94% of C is converted to U during bisulfite conversion. To experimentally test the effect of RP-H, we compared snmC-seq libraries generated from medial frontal gyrus as well as snmC-seq + SAP and snmC-seq2 libraries generated from BA10 (Fig.2b and Supplementary Table 1). Sequencing libraries generated using RP-H contain significantly less adapter dimer reads than those generated with RP-N (p=9.2×10^−10^, Fig.2b).

**Figure 2.**
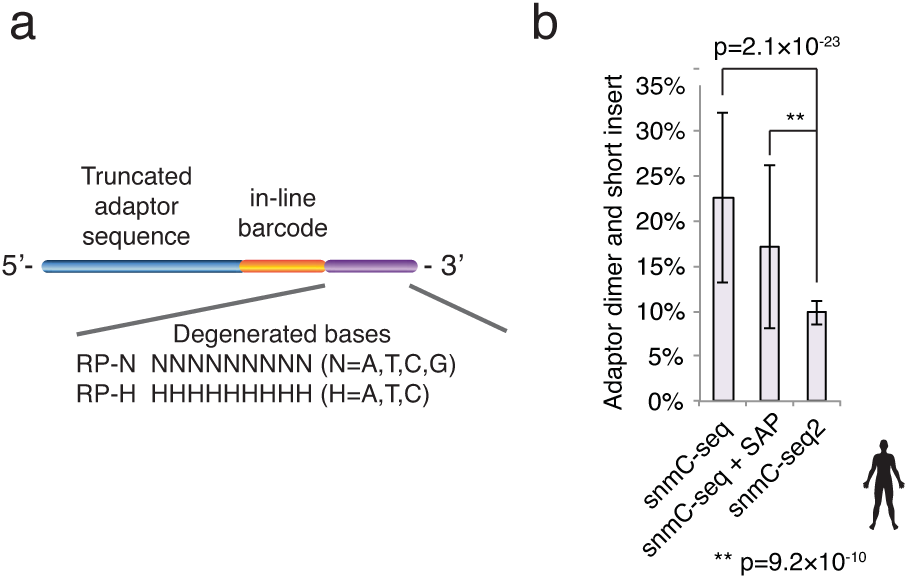
Reduction of adapter dimer and short insert reads by modulating the degenerate 3’ arm of random primers. **(a)** RP-H and RP-H oligos are used for snmC-seq and snmC-seq2, respectively. **(b)** Adapter dimer and short insert reads were significantly reduced in snmC-seq2 libraries.

We have also improved the throughput of single-cell methylome library preparation by developing a 384-well DNA binding plate for cleaning up bisulfite conversion reactions. Compared to the currently available 96-well silica column plates, our newly designed 384-well DNA binding plate provided 33% (p=9.2×10^−10^, t-test) greater library complexity (Supplementary Fig.2a), suggesting more efficient retention of bisulfite-converted DNA from single cells. An advantage of snmC-seq is the multiplexing of 4 samples for 3’-adaptor tagging and all following steps (Step 4 in Fig.1a) ^5^. Individual samples were indexed using “in-line” barcodes located upstream of 3’ degenerate arm of random primers (Fig. 2a). Sample multiplexing was enhanced in snmC-seq2 by reducing the reaction volume of random-primed DNA synthesis, which allowed 8 reactions to be combined for 3’-adaptor tagging. A fully automated robotic protocol has been developed for snmC-seq2, which enabled the preparation of 3,072 single nucleus methylome libraries per experiment. We have developed 768 pairs of library amplification primers containing dual unique indices to avoid “index-hopping” on patterned flowcells (Supplementary Table 2) ^9^. Together with 8 “in-line” barcodes, the primer set allow the multiplexing of 6,144 single-cells for pooled sequencing. Robotic scripts are provided in Supplementary File 2. We tested the snmC-seq2 method by generating single nucleus methylomes from mouse primary motor cortex (MOp, Supplementary Table 1). Single nucleus methylome libraries generated for MOp contain markedly reduced (6.1±5.2%) adapter dimer and short insert reads, compared to that of snmC-seq libraries (29.2±16.5%, Table 1)^5^. Reads mapping rate was improved from 52±4% for snmC-seq to 64.7 ± 2.6% for snmC-seq2. In addition, snmC-seq2 shows enhanced library complexity (maximal coverage of 30.8±7.5% of mouse genome) compared to 22.2±5.7% for snmC-seq. We further examined whether replacing RP-N with RP-H affects the coverage uniformity of snmC-seq2 libraries. Consistent with previous reports ^3^, GC-rich regions such as CpG islands (CGI) are enriched in methylomes generated with PBAT-derived strategies including snmC-seq (Supplementary Fig.2b). Enrichment of CGI was reduced in snmC-seq2 libraries, which also showed more uniform genome coverage for 1kb and 10kb genomic bins than snmC-seq, comparable to traditional MethylC-seq (Supplementary Fig. 2c).

To rigorously evaluate the methylome profiles generated by snmC-seq2, we compared mouse frontal cortex profiled with snmC-seq^5^, MOp analyzed with snmC-seq2 along with cortical excitatory neurons analyzed with MethylC-seq ^10^. Visual examination revealed consistent mCG and mCH levels across all three methods, and uniform snmC-seq2 read coverage in the genomic region surrounding Neurog2 locus (Fig.3a). Further analyses revealed strong correlations between snmC-seq2 and snmC-seq for the quantification of mCH at 1kb genomic bins, (r = 0.959) and mCG at differentially methylated regions (DMRs, r = 0.971) identified in our previous study across all mouse cortical neuron types ^5^.

**Figure 3.**
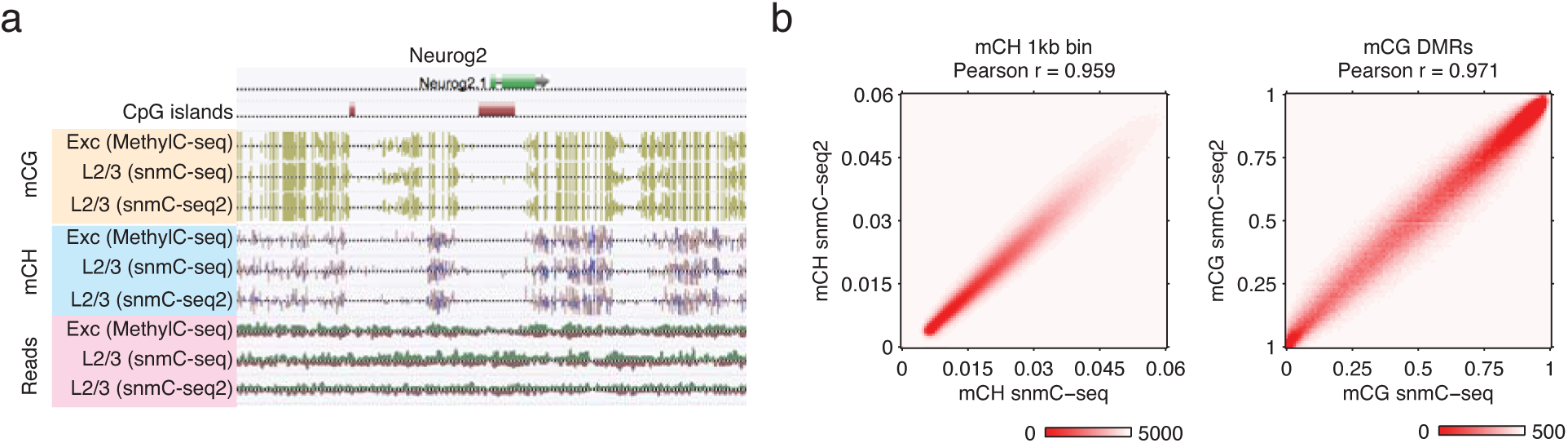
snmC-seq2, snmC-seq and MethylC-seq generate consistent methylome profiles. (**a**) Comparison of CG, CH methylation and read coverage in the region surrounding Neurog2. L2/3 indicates combined methylome profiles of Layer 2/3 excitatory neuron clusters. (**b**) Correlation of CH and CG methylation by snmC-seq2 and snmC-seq were compared for 1kb genomic bins and CG-DMR regions, respectively.

In this study, we introduced snmC-seq2, a method for single nucleus methylome library preparation providing significant improvements in virtually all pertinent aspects including read mapping, amount of artifactual reads, throughput, library complexity and coverage uniformity. The development of snmC-seq2 significantly narrows the gap of sequencing library quality between single-cell and bulk methylomes and provides a compelling strategy to generate large-scale single-cell methylome datasets for the survey of epigenomic diversity across human body cell types and dynamic cellular processes.

## Acknowledgement

This work is supported by NIH BRAIN Initiative grants 5U01MH105985 (J.R.E and M.M.B), 5R21MH112161 (J.R.E and M.M.B), U19MH114831 (J.R.E) and 1R21HG009274 (J.R.E). J.R.E. is an investigator of the Howard Hughes Medical Institute. Postmortem human brain tissues were obtained from the NIH NeuroBioBank at The University of Miami Brain Endowment Bank, and we thank the donors and their families for their invaluable donations for the advancement of science.

## Methods

A detailed bench protocol is provided in Supplementary File 1.

## Animal protocols, dissection, nuclei extraction and FACS isolation of single nuclei

Male C57Bl/6J mice were purchased from Jackson laboratories at 8 weeks of age and maintained in our facility for 48h before dissection. Animals were maintained in the Salk animal barrier facility on 12h dark-light cycles with food ad-libitum. Brains were extracted and sliced coronally at 600 µm from the frontal pole across the whole brain in ice cold dissection buffer containing 2.5mM KCl, 0.5mM CaCl_2_, 7mM MgCl_2_, 1.25mM NaH_2_PO_4_, 110mM sucrose, 10mM glucose and 25mM NaHCO_3_. The solution was kept ice-cold and bubbled with 95%O_2_/5%CO_2_ for at least 15 min before starting the slicing procedure. Slices were kept in 12-well plates containing ice-cold dissection buffer until dissection (normally for approximately 20 min) using a SZX16 Olympus microscope equipped with an SDF PLAPO 1XPF objective. The MOp region was manually dissected from two 600 µm consecutive coronal sections at the following Bregma coordinates: AP 2.36 to 1.16 mm; L 1.00 to 2.75 mm. Only cortical tissue was dissected and snap frozen in dry ice. Tissue from 15 males for each of two biological replicas was processed for nuclei preparation as previously described ^5,8^. Single nuclei were sorted into 384-well PCR plates containing 2µL Digestion Buffer per well. 20mL Digestion buffer contains 10mL M-digestion buffer (2X, Zymo D5021-9), 1ml Proteinase K (20mg, Zymo D3001-2-20), 9mL water and 10µL unmethylated lambda DNA (100pg/µL, Promega, D1521).

## Human brain tissues

Postmortem human brain biospecimens were obtained from brain donors of the University of Miami Brain Endowment Bank. snmC-seq with Shrimp Alkaline Phosphatase treatment (snmC-seq + SAP) was applied to prefrontal cortex (Brodmann Area 10, BA10) tissues obtained from a 58-year-old Caucasian male with a postmortem interval (PMI) = 23.4 hours. snmC-seq2 was applied to BA10 cortical tissue of a 25-year-old Caucasian male with a PMI = 20.8 hours. Both donors were confirmed to have no known neuropathological changes by an neuropathologist.

## Bisulfite conversion

15µL CT conversion reagent (Zymo D5003-1) was added to each well of the 384-well plate. The plates were incubated for 98°C for 8 min, then 64°C for 3.5 hours followed by 4°C. All subsequent centrifugation steps were performed for 5 minutes at 5000xg. 80µL M-Binding Buffer (Zymo D5006-3) was added to Zymo-Spin 384 Well Plate (Zymo C2012), and bisulfite converted samples were transferred to these plates and mixed by pipetting, and centrifuged. 100µL M-Wash Buffer (Zymo D5040-4) was added and the plates were centrifuged. 50µL M-Desulphonation Buffer (Zymo D5040-5) was added and the plates were incubated at room temperature for 15 minutes, then centrifuged. The plates were washed twice with 100uL M-Wash Buffer, then eluted into a clean 384-well plate with 7µL EB Buffer (Qiagen 19086) containing 500nM RP-N or RP-H primers.

## snmC-seq2 library preparation

Bisulfite-converted samples were denatured at 95°C for 3 minutes, then placed on ice for 2 minutes. 5µL Random Priming master mix [1µL 10x Blue Buffer (Enzymatics B0110), 0.25µL Klenow Exo-(50U/µL, Enzymatics, P7010-HC-L), 0.5µL dNTP (10mM each, NEB N0447L), 3.25µL water] was added and incubated at 4°C for 5 min, 25°C for 5 min, and 37°C for 60 min followed by 4°C. 1.5µL enzyme mix containing 1 µL Exonuclease 1 (20U/µL, Enzymatics X8010L) and 0.5 µL rSAP (1U/µL, NEB M0371L) was added, then incubated at 37°C for 30 min followed by 4°C. 0.8x SPRI beads were added, mixed and incubated for 5 minutes at room temperature to allow DNA to bind. The beads were washed 3 times with 80% ethanol and eluted in 10µL EB buffer (Qiagen 19086). The samples were denatured again at 95°C for 3 minutes, and 10.5µL Adaptase master mix [2µL G1, 2µL G2, 1.25µL G3, 0.5µL G4 and 0.5µL G5 (Swift Biosciences 33096)] was added and incubated at 37°C for 30 minutes, then 95°C for 2 minutes. 25µL 2x KAPA HiFi HotStart ReadyMix (Kapa, KK2602) and 5µL custom indexing primer mix (6µM P5, 10µM P7) were added. The PCR was programmed as follows: 1) 95°C 2 min, 2) 98°C 30 sec, 3) 98°C 15 sec, 4) 64°C for 30 sec, 5) 72°C for 2 min, 6) 72°C 5 min, 7) 4°C hold. Repeat steps 3-5 for 15 total cycles. PCR reactions were cleaned with 0.8X SPRI beads for three rounds. Library concentration was determined with Qubit™ dsDNA BR Assay Kit (ThermoFisher Q32853). Libraries were sequenced using Illumina Novaseq instrument.

## Bioinformatics

Sequencing reads mapping, quality filtering and the summary of DNA methylation level for each cytosine was performed as previously described with minor modifications^5^. Non-clonal mapped reads were filtered for MAPQ > 10 using *samtools view -bq10* option. MethylC-seq data generated from pan-excitatory mouse cortical neurons were used for the comparison with snmC-seq and snmC-seq2 datasets (GSM1541958, GSM1541959)^10^. Methylome profiles generated by snmC-seq or snmC-seq2 for Layer 2/3 excitatory neurons were aggregated for the comparison with MethylC-seq data. Layer 2/3 excitatory neurons in the snmC-seq dataset was annotated in our previous study^5^.

To identify Layer 2/3 excitatory neurons in snmC-seq2 dataset generated for mouse primary motor cortex, we computed CH methylation ratio of each non-overlapping 100kb bins across the genome, defined as the number of methylated basecalls divided by the number of total basecalls in the bin with CH context. We only included 18893 bins with >100 total basecalls in more than 97.5% cells in the downstream analysis. Bin-level methylation ratios were divided by the global mCH level of each cell analyzed. We performed principal component analysis on the cell-by-bin methylation ratio matrix and use the top 150 principal components for hierarchical clustering. The expression level of a gene is known to show anti-correlation with gene body mCH level^8,10^, we therefore assigned the clusters to putatitive cell types based on the gene body mCH level of the established marker genes. Layer 2/3 excitatory neurons were identified as the cell cluster with low mCH level at Cux1 (marker of Layer 2/3 and L4) and Satb2 (marker of excitatory neurons) and high mCH level at Rorb (marker of L4 and L5a). The identified L23 neurons are robust to different number of principal components used for clustering.

Preseq was used to estimate library complexity using forward reads with Preseq gc_extrap function with options -e 5e+09 -s 1e+07^11^. Library complexity values shown in this study were estimated for the sequencing depth of 50 million read pairs.

## Data Availability

Raw data and processed data are available from NCBI GEO accession GSE112471. The comparison of DNA methylome generated from mouse cortical excitatory neurons MethylC-seq, snmC-seq and snmC-seq2 can be visualized at http://neomorph.salk.edu/snmC-seq2.php.

## Supplementary figure legends

**Supplementary Figure 1.**
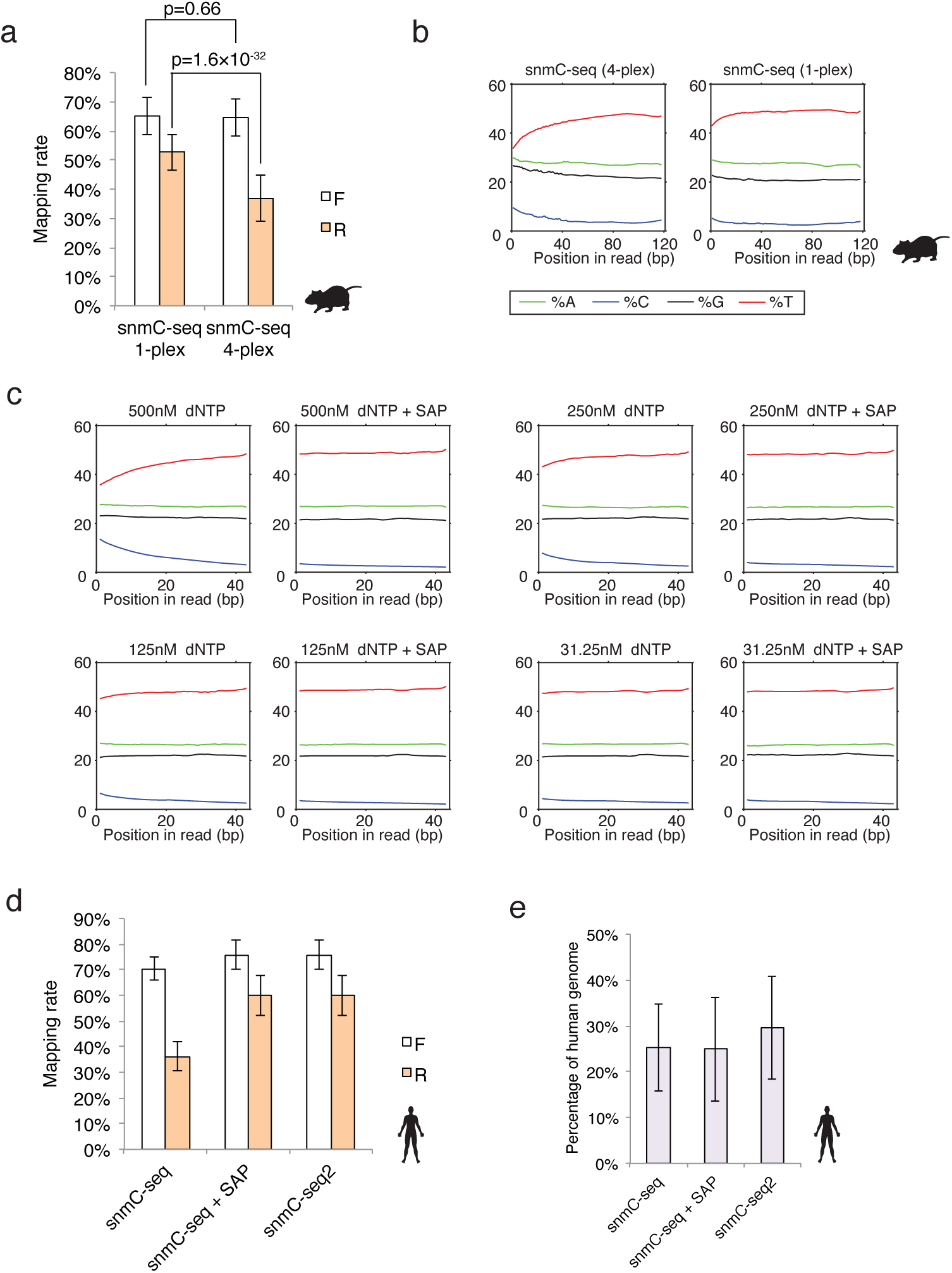
(a) Multiplexed (4-plex) snmC-seq shows reduced reverse reads (R) mapping compared to non-multi-plexed snmC-seq (1-plex). (b) Multiplexed snmC-seq shows aberrant reverse reads base composition. (c) Reducing dNTP concentration in random-primed DNA synthesis or treatment with Shrimp Alkaline Phosphatase (SAP) suppresses the aberrant reverse reads base composition. (d) SAP treatment increases the mapping rate of reverse reads for single-cell methylomes generated from human cortical tissues. (e) Comparison of library complexity for libraries generated with snmC-seq, snmC-seq with SAP treatment (snmC-seq + SAP) and snmC-seq2 using single nuclei isolated from human cortical tissues.

**Supplementary Figure 2.**
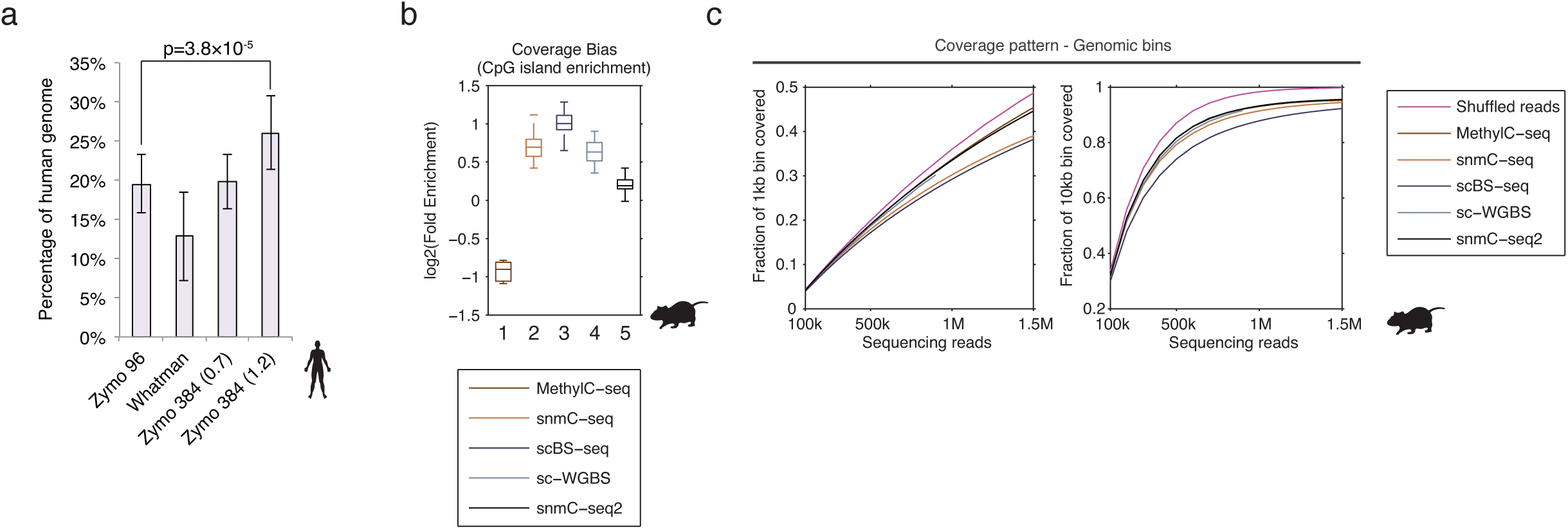
(a) Library complexity for single cell methylomes generated using different DNA binding column plates. Zymo 96 - Zymo-Spin I-96 Binding Plates (Zymo C2004). Whatman - 384 Wells Unifilter Microplate (Whatman 7700-2110). Zymo 384 (0.7) - 384-well DNA binding column plates with glass fiber pore size 0.7 µm. Zymo 384 (1.2) - 384-well DNA binding column plates with glass fiber pore size 1.2 µm. (b) Enrichment of CpG islands in DNA methylome generated by traditional MethylC-seq1, snmC-seq2, scBS-seq3, sc-WGBS4 and snmC-seq2. (c) Fraction of 1kb and 10kb non-overlapping bins covered by single cell methylome reads as a function of the sequencing depth.

## Supplementary File 1 - snmC-seq2 protocol

Protocol to bisulfite convert and generate 3,072 single-cell methylome libraries in one experiment (8x 384-well plates) from mouse or human frozen brain tissue.

## Materials

### General reagents

- FACS sorted single-nuclei into 384 well PCR plate
- HyClone™ HyPure™ Molecular Biology Grade (MB) Water (GE Life Sci. cat. no. SH30538.03)
- 200-Proof (100%) Ethanol (Koptec cat. no. V1001)

### Collection of single nuclei by Fluorescence-activated Cell Sorting (FACS)

- M-Digestion Buffer (Zymo cat. no. D5021-9)
- Proteinase K (Zymo cat. no. D3001-2-D)
- Proteinase K Storage Buffer (Zymo cat. no. D3001-2-B)
- Unmethylated Lambda DNA (Promega cat. no. D1521, 100pg/µL)

### Bisulfite conversion

- CT Conversion Reagent (Zymo cat. no. D5003-1)
- M-Solubilization Buffer (Zymo cat. no. D5021-7)
- M-Dilution Buffer-Gold (Zymo cat. no. D5006-2)
- M-Reaction Buffer (Zymo cat. no. D5021-8)
- M-Binding Buffer (Zymo cat. no. D5021-7)
- M-Wash Buffer (Zymo cat. no. D5040-4)
- M-Desulphonation Buffer (Zymo cat. no. D5040-5)
- M-Elution Buffer (Zymo cat. no. D5007-6)

### Random primers

- HPLC purified Random primers were ordered from Integrated DNA Technologies (IDT) P5L_AD002_H /5SpC3/TTCCCTACACGACGCTCTTCCGATCTCGATGT(H1:33340033)( H1)(H1)(H1)(H1)(H1)(H1)(H1)(H1) P5L_AD006_H /5SpC3/TTCCCTACACGACGCTCTTCCGATCTGCCAAT(H1:33340033)( H1)(H1)(H1)(H1)(H1)(H1)(H1)(H1) P5L_AD008_H /5SpC3/TTCCCTACACGACGCTCTTCCGATCTACTTGA(H1:33340033)( H1)(H1)(H1)(H1)(H1)(H1)(H1)(H1) P5L_AD010_H /5SpC3/TTCCCTACACGACGCTCTTCCGATCTTAGCTT(H1:33340033)( H1)(H1)(H1)(H1)(H1)(H1)(H1)(H1) P5L_AD001_H /5SpC3/TTCCCTACACGACGCTCTTCCGATCTATCACG(H1:33340033)( H1)(H1)(H1)(H1)(H1)(H1)(H1)(H1) P5L_AD004_H /5SpC3/TTCCCTACACGACGCTCTTCCGATCTTGACCA(H1:33340033)( H1)(H1)(H1)(H1)(H1)(H1)(H1)(H1) P5L_AD007_H /5SpC3/TTCCCTACACGACGCTCTTCCGATCTCAGATC(H1:33340033)( H1)(H1)(H1)(H1)(H1)(H1)(H1)(H1) P5L_AD012_H /5SpC3/TTCCCTACACGACGCTCTTCCGATCTCTTGTA(H1:33340033)( H1)(H1)(H1)(H1)(H1)(H1)(H1)(H1)

### Sera-Mag Solid Phase Reversible Immobilization (SPRI) beads

- Sera-Mag SpeedBeads Magnetic Carboxylate Modified (GE Healthcare cat. no. 45152105050250)
- Poly(ethylene glycol) PEG 8000 (Sigma cat no. 89510-250G-F)
- TE buffer pH=8.0 (Ambion cat. no. AM9858)
- 5M NaCl
- 1M Tris-HCl pH=8.0
- 0.5M EDTA pH=8.0
- AMPure XP beads (Beckman Coulter cat no. A63881)
- 100 bp DNA ladder (New England Biolabs cat no. N3231L)

### snmC-seq2 library preparation

- Blue Buffer (10x) (Enzymatics cat. no. P7010-HC-L)
- Klenow Exo-(50U/µL) (Enzymatics cat. no. P7010-HC-L)
- Deoxynucleotide Solution Mix (10mM each dNTP) (NEB cat. no. N0447L)
- Exonuclease I (20U/µL) (Enzymatics cat. no. X8010L)
- Shrimp Alkaline Phosphatase (rSAP) (1U/µL) (NEB cat. no. M0371L)
- Sera-Mag SPRI beads
- M-Elution Buffer (Zymo cat. no. 5007-6)
- EB buffer (Qiagen cat. no. 19086)
- Accel-NGS^®^ Adaptase™ Module (Swift Bio cat. no. 330384)
- P5 Indexing Primer (IDT custom DNA oligo, standard desalted)
- P7 Indexing Primer (IDT custom DNA oligo, standard desalted)
- KAPA HiFi HS RM (Kapa cat. no. KK2602)
- Qubit dsDNA BR Assay Kit (Thermo Fisher cat. no. Q32850)

### Equipment

- 384-Well Hardshell PCR Plate Clear, 20 Pcs/pk (Thermo Fisher cat. no. 4483285)
- 96-Well Hardshell PCR Plate GPLE, 20 Pcs/pk (Thermo Fisher cat. no. 4483348)
- Zymo-Spin 384 Well Plate, 2 pack (Zymo cat. no. C2012)
- 2.0mL 96-well Deep Well Polypropylene Plate, Sterilized (USA-SCI. cat. no. 1896-2110)
- Reservoir Single Well 96 Bottom High Profile, Clear (Axygen cat. no. RES-SW96-HP)
- Reservoir Single Well 96 Bottom Low Profile, Clear (Axygen cat. no. RES-SW96-LP)
- 15mL Centrifuge Tubes (Olympus cat. no. 28-103)
- 50mL Centrifuge Tubes (Olympus cat. no. 28-106)
- 1.5µL Eppendorf Tubes (Thermo Fisher cat. no. 02-681-320)
- 300µL 8-Strip Tubes
- Microamp Clear Adhesive Film, 100pc (Thermo Fisher cat. no. 4306311)
- Microporous Film, -20C to 80C, 50p, sterile (USA-SCI. cat. no. 2920-1010)
- Speedball Deluxe Soft Rubber Brayer, 4 inches (Statesville N.C.)
- 37°C Incubator
- 384-well and 96-well Compatible Thermocycler
- DynaMag™-96 Side Magnet (Thermo Fisher cat. no. 12331D)
- DynaMag™-2 Magnet (Thermo Fisher cat. no. 12321D)
- Allegra 25R Centrifuge with S5700 2×96 Swinging Bucket Rotor (Beckman Coulter cat. no. 368954)
  - All centrifuge steps at 5000x*g* for 5 minutes at room temperature unless otherwise specified
- (Optional) Axygen DITI 50µL STE.FIL. Robotics Tips (Thermo Fisher cat. no. EVF-50-RS)
- (Optional) Axygen DITI 180µL STE.FIL. Robotics Tips (Thermo Fisher cat. no. EVF-180-RS)
- (Optional) TECAN Freedom Evo 100 Base Unit with MultiChannel™ Arm MCA 96
- (Optional/Required for TECAN use) Collection Spacer/Adapter, Type 2, 26.5mm, Te-VacS (Tecan US Inc. cat. no. 10760662)

## Reagent Setup

### Ethanol, 80%

To 40mL of 200-proof Ethanol, add 10µL MB water in a 50mL tube. Keep solution sealed when not in use. Prepare ~6 tubes fresh before each library preparation.

### Digestion Buffer Plates (for FACS sorting)

In a 50mL tube, combine 15mL M-Digestion Buffer (incubate for 10 minutes at 37°C to dissolve precipitate) with 14mL MB water. Resuspend one tube Proteinase K with 1mL Proteinase K Resuspension Buffer and add 1mL to the digestion buffer mix. Add 10µL λ DNA to aid determination of bisulfite conversion efficiency. Vortex to mix. Aliquot 2µL per well to 20 384-well plates to use for FACS sorting of stained nuclei (TECAN script). Plates can be stored at 4C.

(Optional) This step can be automated using TECAN Freedom Evo 100 with script - *2uL_Digestion_Buffer*.

### Sera-Mag Solid Phase Reversible Immobilization (SPRI) Beads

1. Mix Sera-Mag SpeedBeads and transfer 1mL to a 1.5ml tube.
2. Place SpeedBeads on a magnetic stand until clears and carefully remove the supernatant. Wash the beads twice with 1ml of TE. For each wash, remove the tube from the magnet and mix by inversions. Resuspend the washed beads in 1ml of TE.
3. Add 9g of PEG 8000 to a new 50ml sterile conical tube.
4. Add 10ml of 5M NaCl to the 50ml tube.
5. Add 500µl of 1M Tris-HCl pH=8.0 and 100ul of 0.5M EDTA pH=8.0 to the 50ml tube.
6. Mix until all dissolves into solution.
7. Add 1ml of resuspended SpeedBeads to the 50ml tube and fill the volume with MB water.
8. Test against AMPure XP beads using 100bp DNA ladder.

### CT Conversion Reagent

Add 7.9 mL M-Solubilization Buffer and 3 mL M-Dilution Buffer to a bottle of CT Conversion Reagent. Shake vigorously at room temperature to fully dissolve before adding 1.6 mL M-Reaction Buffer.

### M-Wash Buffer

Add 288mL 200-proof Ethanol to four bottles of M-Wash buffer. Invert to mix. Make fresh each time. Extra buffer can be stored at room temperature.

### Random Primer solution

35µL random primer stock (100µM) in 7mL M-Elution buffer (500nM final primer concentration). Aliquot to eight 96-well plates.

### Random Priming Master Mix

Combine 3600µL Blue Buffer (10x), 900µL Enzymatics Klenow exo-(50U/µL), 1800µL dNTP (10mM each), and 11700µL MB water in a 15mL tube and vortex to mix. Aliquot 180µL of master mix into each well of one 96-well plate. Keep sealed on ice until use. Do not store.

### Exo/rSAP Master Mix

Combine 3600µL Exonuclease 1 (20U/µL) with 1800µL rSAP (1U/µL) in a 15mL tube and vortex to mix. Aliquot 53µL master mix to each well on one 96-well plate. Keep sealed on ice until use. Do not store.

### Adaptase Master Mix (using Accel-NGS^®^ Adaptase™ Module)

In a 15mL tube, to 1912.5µL M-Elution buffer, add 900µL Buffer G1, 900µL Reagent G2, 562.5µL Reagent G3, 225µL Enzyme G4, and 225µL Enzyme G5. Aliquot 295µL to 2 sets of 8-strip tubes (16 tubes total). Keep sealed on ice until use. Do not store.

### PCR Primer Mix

Sequences of indexing primers with unique dual barcodes are provided in Supplementary Table 2. Each PCR Primer Mix contains a P5L indexing primer (600 nM) and a P7L indexing primer (1 µM).

## Procedure

### Bisulfite Conversion

Timing ~ 5h

1. Add 15µL CT conversion reagent to each well of 384-well plate. Pipette up and down for 8 times to mix the sample.

(Optional) This step can be automated using TECAN Freedom Evo 100 with script - *15uL_Conversion_Buffer*

2. Seal the plate with adhesive film and quick spin for 10s at 2,000x*g* at room temperature. Place the plate in a thermocycler and run the following program:

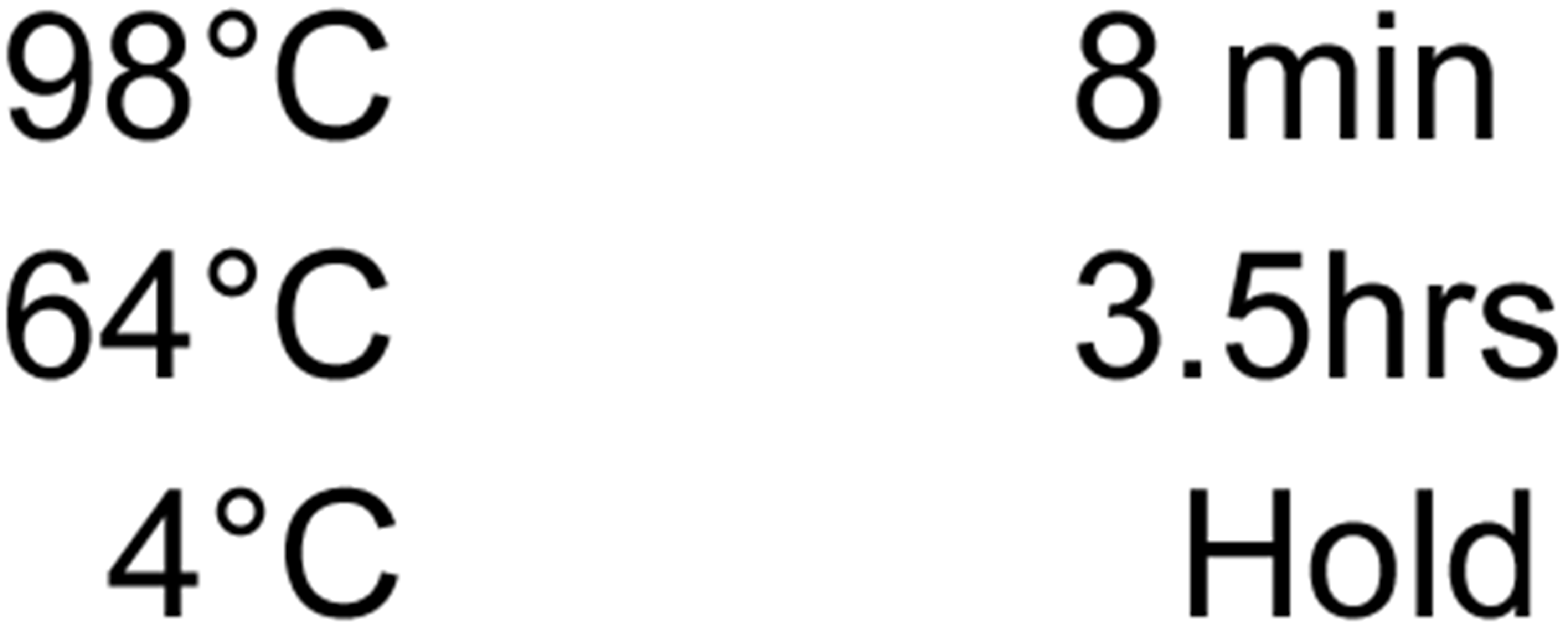

3. Load 80µL M-Binding buffer to each well of 384-Well DNA Binding Plate.

(Optional) This step can be automated using TECAN Freedom Evo 100 with script - *80uL_Binding Buffer*

4. Transfer bisulfite conversion reactions to Zymo-Spin 384 Well DNA Binding Plate. Pipette up and down for 8 times to mix the sample. Place the 384-Well DNA Binding Plate on a 2.0mL 96-well Deep Well Plate and centrifuge for 5 min at 5,000g. Discard the flow through by decanting.

(Optional) This step can be automated using TECAN Freedom Evo 100 with script - *17uL_Conversion_Rxn_To_Binding_Buffer*.

5. Add 100µL M-Wash Buffer to each well of 384-Well DNA Binding Plate. Centrifuge for 5 min at 5,000g and discard the flow through by decanting.

(Optional) This step can be automated using TECAN Freedom Evo 100 with script - *100uL_M_Wash_Buffer*.

6. Add 50µL M-Desulphonation Buffer to each well of 384-Well DNA Binding Plate. Incubate at room temperature for 15 min. Centrifuge for 5 min at 5,000g and discard the flow through by decanting.

(Optional) This step can be automated using TECAN Freedom Evo 100 with script - *50uL_Desulphonation_Buffer*.

7. Add 100µL M-Wash Buffer to each well of 384-Well DNA Binding Plate. Centrifuge for 5 min at 5,000g and discard the flow through by decanting.

(Optional) This step can be automated using TECAN Freedom Evo 100 with script - *100uL_M_Wash_Buffer*.

8. Repeat step 7.

9. Place 384-Well DNA Binding Plate on new 384-well PCR plate. Add 7µL Random Primer Solution. Each quadrant of 384-well plate is barcoded with a distinct indexed random primer (Fig. 1). Every two 384-well plates receive a complete set of all eight indexed random primers. Incubate for 5 min at room temperature. Centrifuge for 5 min at 5,000g and discard the 384-Well DNA Binding Plate.

(Optional) This step can be automated using TECAN Freedom Evo 100 with script - *7uL_Primer_Elution*.

**Figure 1.**
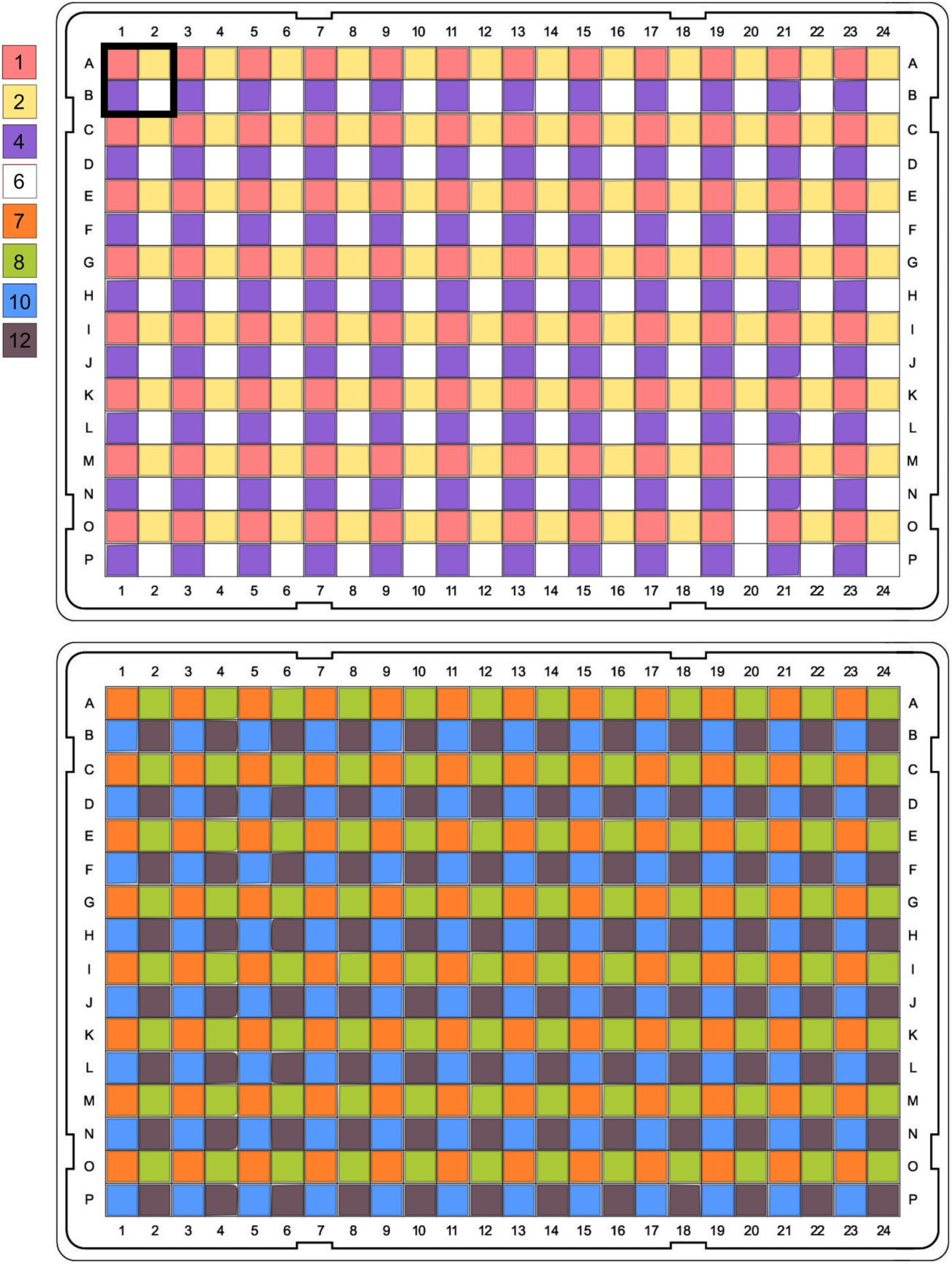
Schematics of single-cell barcoding with indexed random primers. Using a TECAN Freedom Evo 100 with MultiChannel Arm MCA 96, each indexed random primer is added into a quadrant of 384-well DNA Binding Plate. The color of each quadrant indicates the random primer barcode.

10. Seal the plate with adhesive film and store at -20°C for up to 1 week.

### Random-primed DNA synthesis

Timing ~2h

1. Denature the samples by placing 384-well PCR plate on a thermocycler and run the following program.

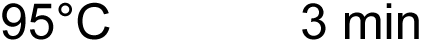 Immediately place the plate on ice for 2 minutes.
2. Add 5µL Random Priming Master Mix to each well of the 384-well PCR plate. Vortex and quick spin for 10s at 2,000x*g*. (Optional) This step can be automated using TECAN Freedom Evo 100 with script - *5uL Random_Priming_Mix*.
3. Place the plate in a thermocycler and run the following program:

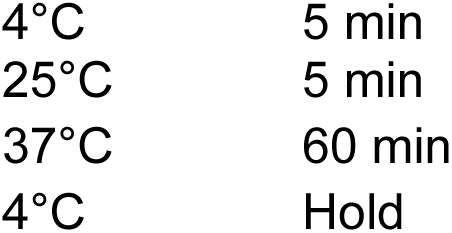

### Inactivation of free primers & dNTP

4. Add 1.5µL Exo/rSAP Master Mix to each well of the 384-well PCR plate. Vortex to mix the samples and quick spin for 10s at 2,000x*g*.

(Optional) This step can be automated using TECAN Freedom Evo 100 with script - *1uL_Exo_rSAP_MM*.

5. Place the plate in a thermocycler and run the following program

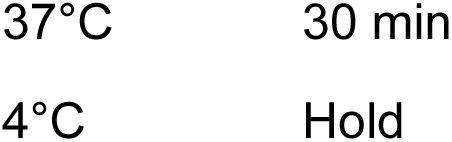

### Sample clean-up

6. Add 73.6µL (0.8x) SPRI Beads to each well of four clean 96-well PCR plates.

(Optional) This step can be automated using TECAN Freedom Evo 100 with script - *BP_Add_To_Clean_96well*.

7. Transfer samples from two 384-well plates to each 96-well plate. The eight quadrants (from two 384-well plates) barcoded with distinct indexed random primers are combined. Mix the samples by vortexing and incubate for 5 minutes at room temperature, then quick spin for 10s at 2,000x*g*.

(Optional) This step can be automated using TECAN Freedom Evo 100 with script - *BP_Transfer_Lib_384well_To_96well*.

8. Place 96-well plates on DynaMag™-96 Side Magnet, let stand until solution in wells is clear of beads (~5 minutes). Remove supernatant and wash beads 3x with 150µL fresh 80% EtOH. Remove all EtOH, remove plates from magnet, and let beads dry at room temperature. DO NOT overdry beads.

(Optional) This step can be automated using TECAN Freedom Evo 100 with script - *BP_2Plate_150uL_EtOH_Wash*.

9. Add 10µL EB buffer and resuspend beads by pipet. Vortex and incubate for 5 minutes at room temperature, then quick spin for 10s at 2,000x*g.* Place back on magnet and let stand until solution is clear of beads (~5 minutes).

10. Remove 10uL supernatant to a clean 96-well PCR plate.

(Optional) This step can be automated using TECAN Freedom Evo 100 with script - *10ul_Pre_Adaptase_Transfer*.

### Adaptase reaction

11. Denature the samples by placing 96-well plates on a thermocycler and run the following program.

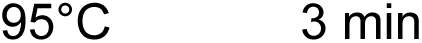

Immediately place the plate on ice for 2 minutes.

12. Add 10.5µL Adaptase Master Mix to each well of the 96-well PCR plate. Vortex and quick spin for 10s at 2,000x*g*.

13. Place the plate in a thermocycler and run the following program:

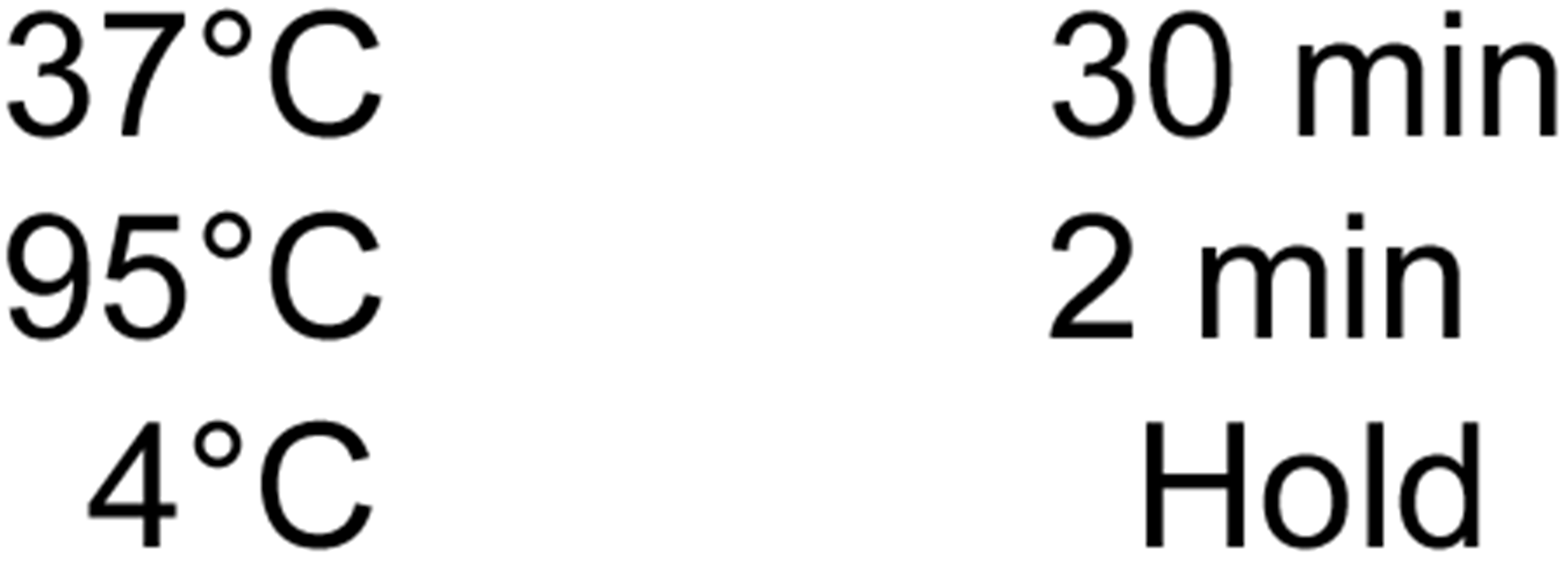

### Library amplification

14. Add 5uL PCR Primer Mix.

(Optional) This step can be automated using TECAN Freedom Evo 100 with script - *5uL_PCR_Primer*.

15. Add 25µL 2x KAPA HiFi Mix. Vortex and quick spin for 10s at 2,000x*g*.

16. Place the plate in a thermocycler and run the following program:

|  |  |  |
| --- | --- | --- |
| a. | 95°C | 2 min |
| b. | 98°C | 30s |
| c. | 98°C | 15s |
| d. | 64°C | 30s |
| e. | 72°C | 2 min |
| Go to step c 14 times |  |  |
| f. | 72°C | 5 min |
| g. | 4°C | Hold |

### Library clean-up

17. Sample cleanup. Add 40µL (0.8x) SPRI Beads to each well of four 96-well PCR plates. Transfer contents of one 96-well plate to another to combine four 96-well plates to two 96-well plates. Vortex and incubate for 5 minutes at room temperature, then quick spin for 10s at 2,000x*g*.

(Optional) This step can be automated using TECAN Freedom Evo 100 with script - *BP_Add_Beads_And_Combine_96well*.

18. Place 96-well plates on DynaMag™-96 Side Magnet, let stand until solution in wells is clear of beads (~5 minutes). Remove supernatant and wash beads 2x with 150µL fresh 80% EtOH. Remove all EtOH, remove plate from magnet and let beads dry at room temperature. DO NOT overdry beads.

(Optional) This step can be automated using TECAN Freedom Evo 100 with script - *BP_2Plate_150uL_EtOH_Wash (stop script after two wash steps)*.

19. Add 25µL EB and resuspend beads by pipet. Vortex and incubate for 5 minutes at room temperature, then quick spin for 10s at 2,000x*g.* Place back on magnet and let stand until solution is clear of beads (~5 minutes). Remove 25uL supernatant to a clean 96-well PCR plate.

20. Add 20µL (0.8x) SPRI Beads to each well of two 96-well PCR plates. Transfer contents of one 96-well plate to another to combine two 96-well plates to one 96-well plate. Vortex and incubate for 5 minutes at room temperature, then quick spin for 10s at 2,000x*g*.

(Optional) This step can be automated using TECAN Freedom Evo 100 with script - *BP_Add_Beads_And_Combine_96well*.

21. Place 96-well plate on DynaMag™-96 Side Magnet, let stand until solution in wells is clear of beads (~5 minutes). Remove supernatant and wash beads 2x with 150µL fresh 80% EtOH. Remove all EtOH, remove plate from magnet, and let beads dry at room temperature. DO NOT overdry beads.

(Optional) This step can be automated using TECAN Freedom Evo 100 with script - *BP_1Plate_150uL_EtOH_Wash*.

22. Add 20µL EB buffer and resuspend beads by pipet. Vortex and incubate for 5 minutes at room temperature, then quick spin for 10s at 2,000x*g.* Place back on magnet and let stand until solution is clear of beads (~5 minutes). Combine 20µL eluent from all wells in each column of the 96-well plate (8 wells per column, 12 columns) into 12 1.5µL Eppendorf tubes.

23. Add 128µL (0.8x) SPRI Beads to each 1.5µL Eppendorf tube. Pipette to mix and incubate for 5 minutes at room temperature.

24. Place 1.5µL tubes on DynaMag™-2 Magnet, let stand until solution in tubes in clear of beads (~5 minutes). Remove supernatant and wash beads 2x with 200uL fresh 80% EtOH. Remove all EtOH, remove tubes from magnet, and let beads dry at room temperature. DO NOT overdry beads.

25. Add 40µL EB and resuspend beads by pipet. Incubate for 5 minutes at room temperature. Place tubes back on magnet and let stand until solution is clear of beads (~5 minutes). Remove 40uL supernatant to 12 clean 1.5µL Eppendorf tubes.

26. Measure concentration of each 1.5µL Eppendorf tube with Qubit dsDNA BR Assay Kit. Normalize library concentrations and pool for sequencing.

